# Oncogene-driven metabolic regulation of Dihydroceramide Desaturase 1 (DES1) converges on GAPDH in matrix-detached conditions

**DOI:** 10.64898/2026.06.18.733233

**Authors:** Danielle L. Lambadis, Victoria Franzi, Deanna M. Peperno, Ryan W. Linzer, Jonathan Aminov, Hector R. Romero Garcia, Charly N. Campanella, Andrew E. Resnick, Folnetti A. Alvarez, Janet J. Allopenna, Christopher J. Clarke

**Author notes:** Correspondence: Christopher J. Clarke, PhD Associate Professor Stony Brook Cancer Center 1 Lauterbur Drive Stony Brook, NY 11794.

## Abstract

Deregulation of sphingolipid (SL) metabolism is well-established across many cancers, yet the underlying mechanisms that drive changes in SLs are poorly understood. We previously identified dihydroceramide desaturase 1 (DES1) as a downstream target of HER2 and implicated DES1 as a driver of anchorage-independent survival in breast cancer. In this study, we expand on these results to establish the oncogenic PI3K pathway as a driver of post-translational DES1 activity following cell detachment from the extracellular matrix. PI3K activation of DES1 required glucose uptake and metabolism through both glycolysis and the pentose phosphate pathway. However, it did not require glucose flux into the TCA cycle and was independent of antioxidant capacity of the cell. Moreover, Instead, results identify GAPDH – a point of convergence between glycolysis and PPP – as important for oncogene-driven DES1 activity. Overall, this study defines a novel pathway of DES1 regulation and establishes DES1 as a point of crosstalk between glucose and SL metabolic pathways.

---

Alterations in cellular metabolism are now appreciated as a hallmark of cancer cells (1). The best-characterized metabolic adaptation is the Warburg effect, where cancer cells preferentially utilize aerobic glycolysis over oxidative phosphorylation (2). Although initially thought to be a by-product of transformation or a consequence of defective mitochondria, this ultimately led to the discovery that oncogenic signaling directly reprograms cell metabolism. Among the major oncogenic pathways, phosphoinositide 3-kinase (PI3K) signaling plays a central role in metabolic reprogramming (3, 4) and regulates glucose metabolism at multiple levels: enhancing glucose uptake through regulating GLUT proteins (5, 6); increasing activity of glycolytic enzymes via downstream effectors such as AKT, Rac, and mTOR (7–9); and promoting glucose flux into the pentose phosphate pathway (PPP) through regulation of glucose-6-phosphate dehydrogenase (G6PD)(10, 11). Metabolic reprogramming is important for supporting the bioenergetic and biosynthetic demands of rapidly proliferating tumor cells but also enables them to adapt to environmental stressors which can promote more aggressive tumor behavior. For example, PI3K-dependent glucose uptake and flux through the PPP in breast cancer (BC) is important for anchorage-independent survival (AIS) – a key metastasis enabling biology associated with aggressiveness (11). Similarly, analysis of brain metastatic BC tumors showed profound upregulation of the PPP and other glucose metabolic pathways (12). Consequently, a better understanding of the mechanisms by which altered metabolism drive aggressive phenotypes could define new therapeutic targets of interest.

Sphingolipids (SLs) are a family of bioactive lipids involved in many biologies including proliferation, apoptosis, inflammation, and stress responses (13). Cellular SL levels are controlled by an interlinked network of enzymes and, as with other metabolic pathways, the dysregulation of SLs has been reported across multiple cancers (14–17). While this led to increased interest in modulating SL levels for cancer therapy, efforts to translate *in vitro* findings into actionable targets in the SL network have proven difficult. This is in part due to our poor mechanistic understanding of the role of SLs in tumor behavior but could also be a consequence of limited knowledge of the mechanisms that drive alterations in the SL network in cancer. As with other metabolic pathways, evidence has begun to suggest that SL metabolism is directly influenced by oncogenic signaling. For example, studies in chronic myelogenous leukemia found that the oncoprotein BCR-ABL alters SL levels by increased expression of sphingomyelin synthase 1 (18, 19) and sphingosine kinase 1 (SK1)(20) as well as increased phosphorylation and activation of SK1 (21). Similarly, oncogenic K-Ras-G12V and B-Raf-V600E were found to promote SK1 activity and S1P production (22). Adding to this, we recently investigated interactions between oncogenic HER2 (NeuT) and SL metabolism in BC, leading to the identification of dihydroceramide desaturase 1 (DES1) as a key SL enzyme that was both necessary and sufficient for anchorage-independent survival (AIS) in BC (23). Notably, while DES1 activity was profoundly inhibited in breast epithelial cells upon detachment from extracellular matrix (ECM), NeuT maintained DES1 activity in suspension (23). However, while this established DES1 as a HER2-regulated node in the SL network, it raised the question of whether DES1 functions as a broader target of oncogenic signaling. Clearly, a deeper understanding of such interactions would be particularly important for defining the signaling contexts where they may have most relevance as a therapeutic target.

In de novo SL synthesis, DES1 occupies the last and key step converting dihydroceramide (dhCer) into ceramide (Cer). Despite this important position, surprisingly little is known about mechanisms of DES1 regulation. Several stimuli have been found to post-translationally inhibit DES1 activity including hypoxia (24), oxidative stress (25), and high cell density (26) among others. For hypoxia, this was speculated to be due to low oxygen effects on the associated iron molecules at the core of the DES1 enzyme (24) although intriguingly, long-term hypoxia can lead to reduced DES1 expression (27). Effects at high cell density were attributed to increased levels of free thiols (26) while the mechanisms for oxidative stress were less clear although appeared to be indirect (25). Similarly, little is known about mechanisms that promote DES1 activity at any level. Myristic acid was reported to drive DES1 activity by promoting N-myristoylation and causing translocation to the mitochondria (28, 20), while a separate study connected free palmitate and myristic acid with increased DES1 expression in muscle cells (30). Given that DES1 controls the levels of two important bioactive SLs, a better understanding of its regulation could provide broader insight into the deregulation of SLs during cancer and other pathologies.

Here, we have built upon our prior study to further investigate oncogenic regulation of DES1 and to define the key metabolic pathways required to sustain DES1 activity following ECM detachment. We find that both NeuT and PI3K-driven DES1 activity requires glucose uptake and flux through the PPP and glycolysis but is independent of cellular antioxidant capacity and the TCA cycle. Moreover, studies pinpoint GAPDH – a convergent point of the PPP and glycolysis – as key for oncogene-driven DES1 activity. Collectively, these studies further define a novel pathway of DES1 regulation and establishes DES1 as a point of crosstalk between glucose and SL metabolic pathways.

## EXPERIMENTAL PROCEDURES

### Reagents

Standard laboratory reagents were from Millipore Sigma or Fisher Scientific. Drugs used were: WZB117, BAY876, 2-DG, 3-BP, G6PDi-1, Koningic acid (KA). Antibodies for p-Akt (S473), Akt, p110a, p85, PTEN, and GAPDH were from Cell Signaling; DES1/MLD (ab167169) was from Abcam; V5 was from Life Technologies; Actin was from Millipore Sigma. Taqman assays and siRNA were purchased from Life Technologies. Information on antibodies, taqman assays, and siRNA are in supplemental tables.

### Cell Lines

MCF10A (CRL-10317) were obtained from ATCC (Manassas, Virginia, USA), authenticated by STR profiling, and cultured in F12/DMEM supplemented with 5% horse serum (#16050-122, Life Tech), hydrocortisone (0.5mg/ml; H0888, Sigma), insulin (10mg/ml; I1882, Sigma), cholera toxin (100ng/ml, C8052, Sigma), and epidermal growth factor (EGF) (20ng/ml, AF-100-015, Peprotech) as described previously (23, 31). Cells were incubated in a humidified atmosphere at 37C, 5% CO2 and were confirmed to be mycoplasma free every other month using the MycoAlert Kit from Lonza. Generation of MCF10A cell lines stably expressing a constitutively active Her2 (10A-NeuT) was described previously (23) Generation of MCF10A cell lines stably expressing oncogenic PI3K (10A-PI3K-EK, 10A-PI3K-HR), oncogenic K-Ras (10A-KRas-G12V), and active MEK (10A-MEK-DD) is described below.

### Plasmids

Lentiviral plasmids pCMV-VSV-G (Addgene #8454) and pCMV-dR8.2 dvpr (Addgene #8455) were gifts from Bob Weinberg. Gateway plasmids pENTR4-V5 (Addgene #17425), pLenti CMV/TO Puro Empty (Addgene #17482) and pLenti CMV/TO Puro DEST (Addgene #17293) were gifts from Eric Campeau. The pBabe-PI3K-E545K (Addgene #12525) and pBabe-PI3K-H1047R (Addgene #12524) plasmids were gifts from Jean Zhao. The MEK-DD plasmid (Addgene #15268) was a gift from William Hahn. To generate lentiviral plasmids expressing oncogenic PI3K or MEK-DD, the cDNA from appropriate pBabe plasmids was subcloned into pENTR-V5 and introduced into pLenti CMV/TO Puro DEST by a recombinase reaction with LR Clonase II (Life Technologies) according to manufacturer instructions. All generated plasmids were sequenced prior to use.

### Generation of isogenic stable MCF10A cell lines using lentivirus

To make lentiviral particles, 293T cells (2500K, 100mm dish) were co-transfected with target plasmids (or pLenti Puro-empty as control), VSV-G, and dVPR in a 1:1:1: ratio using Lipofectamine 2000 (Life Technologies) in opti-MEM (GIBCO, Life Technologies) according to manufacturer’s protocol. The next day, media was changed and 24h later, an additional 2-3ml media was added. At 72h post transfection, virus-containing media was harvested, filtered (Millipore 0.45mM PVDF), and divided into 1ml aliquots for storage at −80C. For infection, MCF10A cells (100K) were plated in 6-well trays and 24h later, 1ml of viral supernatant was added to 1ml growth media containing polybrene (#TR-1003-G; Millipore Sigma; final concentration 8mg/ml). After 24h, media was replaced for a further 24h before cells were split into 100mm dishes in media containing puromycin (1-2mg/ml; InVivogen, San Diego, CA). After 7 days of selection, cells were cultured in normal growth medium. In all cases, uninfected cells were cultured in parallel and treated with puromycin for selection to confirm efficacy of antibiotic selection. Typically, all uninfected cells were dead within 48-72h after initiation of selection.

### siRNA transfection and harvesting

For experiments comparing ECM attached and -detached conditions, cells were cultured in standard tissue culture dishes/trays for attached (Att) and ultra-low attachment dishes/trays (Corning #CLS3471) for detached conditions (Susp). When inhibitors were used, cells were plated directly into media containing either inhibitor or vehicle (0.1% DMSO). For siRNA treatment, cells were reverse transfected in 100mm dishes (800-1000K) with 20nM target siRNA or AllStars (si-AS) negative control siRNA (#1027281; Qiagen, Hilden, Germany) using Lipofectamine RNAiMAx (#13778-150; Life Technologies) according to manufacturer’s protocol.

After 48h siRNA treatment, cells were plated in suspension. A list of siRNA used is in supplemental table 1. To harvest monolayer cultures, cells were washed in cold PBS and scraped directly in lysis buffer (RIPA for protein, RNA lysis buffer for RNA, extraction solvent for lipids). To harvest suspension cultures, cells were pelleted by centrifugation (3000rpm, 5m, 4C), washed in 2ml cold PBS, and centrifuged again. Subsequent pellets were resuspended in lysis buffer as above according to the analysis being performed.

### Analysis of cellular protein by SDS-PAGE and immunoblot

For preparation of protein lysates, cells were harvested in 150μl RIPA buffer (50mM Tris, 150mM NaCl, 1% Triton X-100, 0.5%, 0.1% sodium dodecyl sulfate) supplemented with protease inhibitor cocktail (#P-8340, Millipore Sigma) and phosphatase inhibitor cocktails 2 and 3 (#P5726 and #P0044, Millipore Sigma). Cells were lysed by tip sonication (2 x 10s bursts on ice) and protein concentration was determined by the Bradford assay (Biorad reagent #50000006; Biorad, Hercules, CA). Lysates were combined with equal volumes of 2x Laemelli buffer (#1610737, Biorad) containing 5% b-mercaptoethanol (#M6250, Millipore Sigma), vortexed for 2-3s and heated to 100C for 10 mins. Protein samples were stored at −20C until subsequent analysis. SDS-PAGE and immunoblot analysis was performed as described previously (23) using equal protein amounts (10-20mg). Information on primary antibodies used is in supplemental table 2.

### In situ assay to measure dihydroceramide desaturase (DES) activity

Measurement of *in situ* DES activity was performed using cell-permeable C12-dhCCPS substrate as described previously (23, 32). Lipids were analyzed by tandem LC/MS mass spectrometry using 10ml of C12-dihydroceramide (#860635) from Avanti (Alabaster, AL) as internal standard. *In situ* DES activity was calculated as the ratio of detected product (C12-CPPS) to detected substrate (C12-dhCCPS).

### Analysis of cellular SLs

Lipids were extracted by addition of 2ml 2:3 70% Isopropanol: Ethyl Acetate. Samples were processed and analyzed by tandem liquid chromatography/mass spectrometry (LC/MS) at the Stony Brook Lipidomics Core as described previously (23, 32). Cellular SL levels were normalized to total lipid phosphate levels, analyzed as described previously (23, 32).

### Quantitative Reverse Transcriptase PCR (qRT-PCR)

Total RNA was extracted using the Purelink RNA kit (#12183-020, Life Technologies) and RNA concentration was verified by nanodrop. 0.5-1mg of RNA was transformed into cDNA using the Superscript III supermix kit for real-time PCR (#11752-250; Life Technologies) according to manufacturer’s protocol and converted cDNA was diluted up to 200-300ml with molecular biology water. For qRT-PCR, reactions were run in triplicate in 96 well plates (VWR #93001-100) with each reaction containing 10ml of 2X iTAQ universal probes mastermix (#172-5134, Biorad), 5ml of cDNA, 1ml of Taqman primer-probe (Life Technologies) and 4ml of molecular biology water. The list of taqman assays used is in supplemental table 3.

### Statistical Analysis

Data were graphed and analyzed using Graphpad Prism. Analysis of two means was conducted with Student’s t-test; analysis of more than two means by one-way ANOVA with appropriate post-test; and comparison of two variables by two-way ANOVA with appropriate post-test. For all statistical tests, a *p*-value of less than 0.05 was considered as threshold sufficient to reject the null hypothesis.

## RESULTS

### Increased PI3K signaling through oncogenes and loss of tumor suppressors drives DES activity following ECM detachment

We recently identified DES1 as a key node in the SL network that was regulated by oncogenic HER2 in ECM-detached conditions, requiring PI3K but not ERK signaling (23). Notably, the PI3K pathway is one of the most activated across multiple cancers (33, 34), occurring in response to activation of upstream receptors, receptor independent activation/mutation of PI3K or other oncogenes, and loss of key tumor suppressors such as PTEN and p85. Thus, we speculated that DES1 was a downstream effector of multiple oncogenic signaling pathways. To explore this, we used lentivirus to generate stable cell lines expressing two common oncogenic PI3K variants (E545K, H1027R), wild-type PI3K (WT) or empty vector as control (Vec). For this, non-tumorigenic MCF10A cells were used as we had previously shown that >99% of the *in situ* DES activity measured was due to DES1 and not the DES2 isoform (Linzer, 2023). Immunoblot analysis confirmed stable expression of constructs and showed significantly increased p-Akt (S473) in 10A-EK and 10A-HR cell lines with minor increases in p-Akt seen with WT PI3K (**Fig. 1A**). Trypan blue exclusion assays confirmed that both oncogenic PI3K variants conferred anchorage-independent survival (AIS)(**Fig. 1B**). *In situ* activity assays showed that both oncogenic PI3K variants significantly increased DES1 activity in ECM-detached conditions compared to both Vec and WT cell lines (**Fig. 1C**) but had no major effect on DES1 protein levels confirming post-translational activation as seen previously (**Fig. 1D**). Consistent with this, endogenous dhCer levels were significantly decreased in ECM-detached 10A-EK and -HR cells compared to 10A-Vec and - WT cells (**Fig. 1E, left**) although Cer levels were not altered (**Fig. 1E, right**). In addition to increased upstream signaling, PI3K pathway activation also occurs through the loss of negative regulators such as PTEN and the regulatory p85 subunit of PI3K. Both proteins are considered tumor suppressors, and their loss has been reported in various cancers (33–36). To determine if DES1 activation is relevant in this context, siRNA was used to knockdown p85 or PTEN in wild-type MCF10A cells prior to Susp culture for 24h. Both siRNA led to a robust and specific knockdown for each protein although surprisingly this only led to modest increases in p-Akt (S473) (**Fig. 1F**). Nonetheless, loss of either protein was sufficient to increase *in situ* DES activity in susp without significant effects on DES1 protein levels (**Fig. 1G**). Collectively, these results support a role for PI3K as a driver of DES1 activity.

**Fig. 1:**
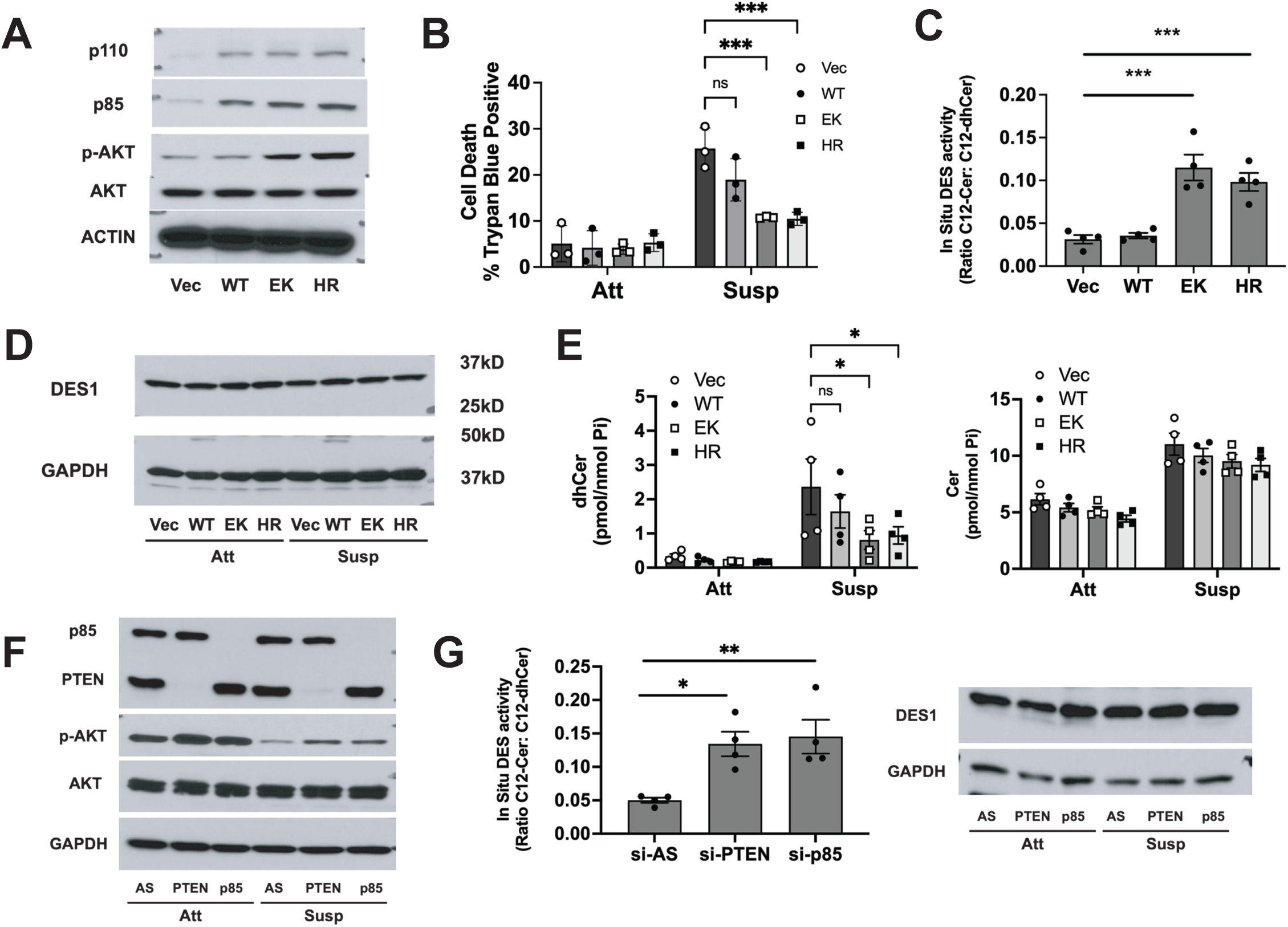
Hyperactivation of the PI3K pathway promotes DES1 activity in ECM-detached conditions. a) Protein was extracted from 10A-Vec, -WT, -EK, and -HR cells and immunoblotted as shown; b) 10A-Vec, -WT, -EK, and -HR were plated in monolayer (Att) or suspension (Susp) for 48h. Cell viability was analyzed by trypan blue exclusion; c) 10A-Vec, -WT, -EK, and -HR cells were plated in susp for 24h and in situ DES activity measured; d) 10A-Vec, -WT, -EK, and -HR cells were plated in Att and Susp conditions for 24h. Protein was extracted and immunoblotted as shown; e) Cells were treated as in ‘d’. Lipids were extracted and analyzed as shown; f) MCF10A cells were treated with si-AS, si-PTEN, or si-p85 for 48h before plating in Att and Susp conditions for 24h. Protein was extracted and immunoblotted as shown; g) 10A cells were treated as in ‘g’, left – *In situ* DES activity was measured, right – protein was extracted and immunoblotted as shown. Data are expressed as mean ± SEM with * p<0.05, ** p<0.02, *** p<0.01 for comparisons shown.

Activation of the PI3K pathway is also key for transformation induced by other oncogenes such as K-Ras. To extend our findings, we utilized MCF10A cells stably expressing the G12V oncogenic K-Ras in MCF10A cells with empty vector as controls (Vec) described previously (37). Immunoblot analysis showed elevations of both p-ERK and p-Akt (S473) consistent with known downstream signaling of K-Ras (**Fig. 2A**), while trypan blue exclusion assays again confirmed the acquisition of AIS by 10A-GV cells (**Fig. 2B**). As with PI3K variants, oncogenic K-Ras led to significant increases in DES1 activity following ECM detachment (**Fig. 2C**). Moreover, 10A-GV cells showed significant lower dhCer levels in ECM-detached conditions compared to control cells (**Fig. 2D, left**) although again, Cer levels were not changed (**Fig. 2D, right**). Finally, pharmacological inhibition of PI3K strongly inhibited K-Ras-GV-induced DES1 activity (**Fig. 2E**) – as seen previously with NeuT (23) with immunoblot analysis confirming inhibition of ERK and Akt (**Fig. 2F**). Intriguingly, and in contrast to NeuT, ERK inhibition led to a modest but significant reduction in K-Ras-G12V-driven DES1 activity (**Fig. 2E)**. While this broadly supports a PI3K-DES1 connection, this does suggest that there may an ERK-dependent arm of DES1 regulation in some contexts. To assess this, MCF10A cells expressing constitutively active MEK (MEK-DD) were generated with immunoblot analysis confirming a modest increase in p-ERK levels in suspension (**Fig. 2G**). However, unlike with oncogenic PI3K, MEK-DD expression was unable to promote DES1 activity in ECM-detached conditions (**Fig. 2H**). Taken together, these results establish DES1 as a central effector of the PI3K pathway and downstream of oncogenic signaling that utilizes the PI3K axis.

**Fig. 2.**
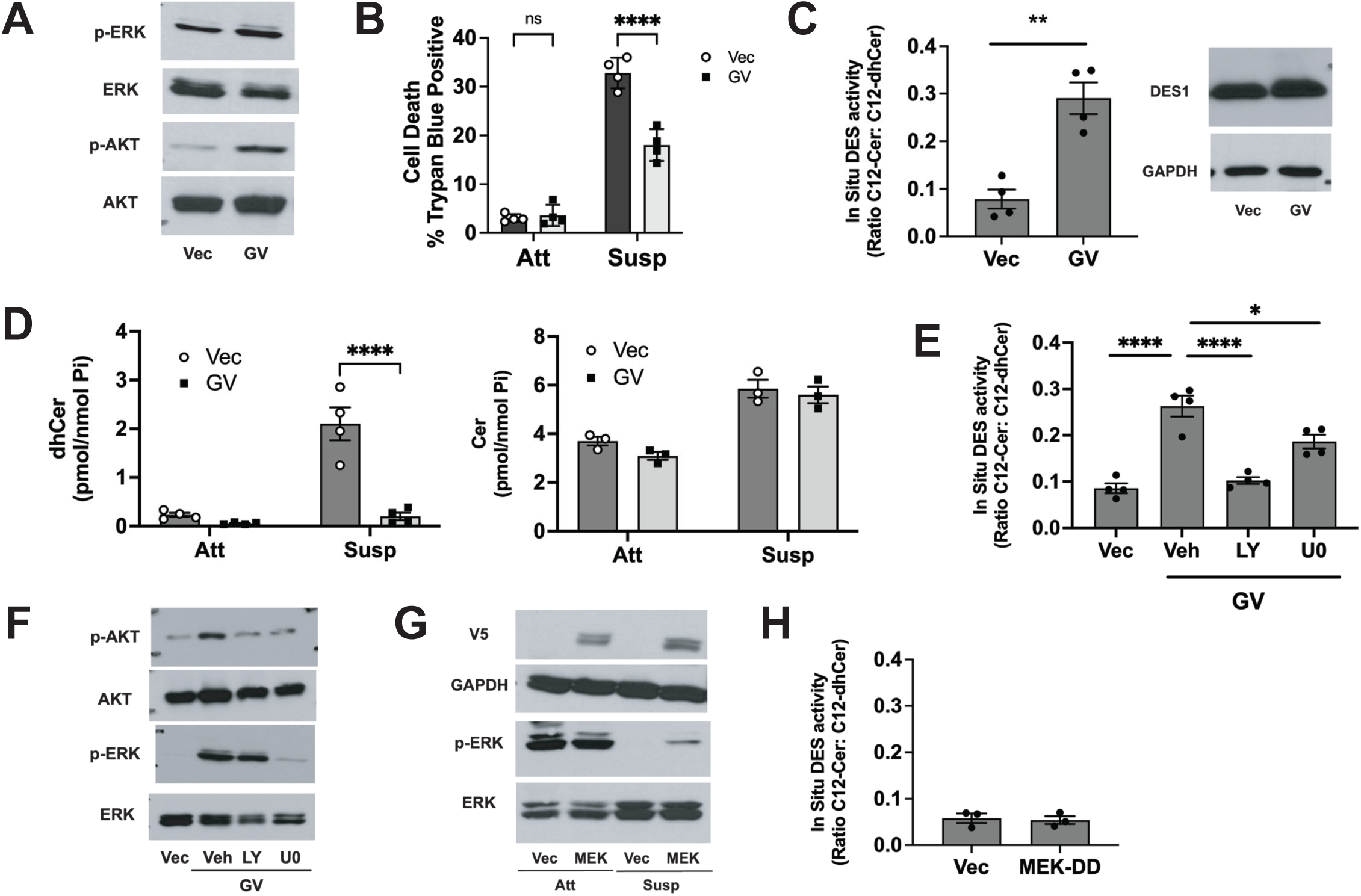
Oncogenic K-Ras drives DES1 activity through PI3K signaling. a) Protein was extracted from 10A-Vec, and K-Ras-GV cells and immunoblotted as shown; b) 10A-Vec, and K-Ras-GV cells were plated in monolayer (Att) or suspension (Susp) for 48h. Cell viability was analyzed by trypan blue exclusion; c) (left) 10A-Vec, and K-Ras-GV cells were plated in susp for 24h and in situ DES activity measured, (right) protein was extracted and immunoblotted as shown; d) 10A-Vec and -GV cells were plated in Att and Susp conditions for 24h. Lipids were extracted and analyzed as shown; e) 10A-Vec and K-Ras-GV were plated in Susp conditions for 24h in the presence of the inhibitors shown. In situ DES activity was measured; f) Cells were plated as in ‘e’ and protein was immunoblotted as shown; g) 10A-Vec and MEK-DD were plated in Att and Susp conditions for 24h and immunoblotted as shown; h) 10A-Vec and MEK-DD were plated in Susp conditions for 24h and in situ DES activity analyzed. Data are expressed as mean ± SEM with * p<0.05, ** p<0.02, **** p<0.001 for comparisons shown.

### Oncogenic PI3K activation of DES1 requires GLUT1-driven glucose uptake and metabolism

PI3K is a well-established regulator of classical glucose metabolic pathways at multiple levels (3–11) and our prior study found that NeuT-driven DES1 activity was dependent on glucose uptake (23). To extend these findings and determine if this was more broadly applicable, oncogenic PI3K cell lines were placed in suspension with the GLUT antagonist WZB117 (20mM)(**Fig. 3A**) with results showing a significant reduction in PI3K-driven DES1 activity for both EK and HR cell lines, but with minimal effect on DES1 protein. Similarly, siRNA knockdown of GLUT1 (**Supp Fig. 1**) prior to suspension culture also led to a significant reduction in PI3K-driven DES1 activity, comparable to the inhibitor effects (**Fig. 3B**). To assess if further glucose metabolism was required for DES activity, 10A-EK and -HR cells were incubated with 2-deoxyglucose (2-DG; 12mM) which blocks onward glucose metabolism (**Fig. 3C**), or 3-bromopyruvate (3-BP; 50mM), a structurally distinct inhibitor of hexokinase (**Fig. 3D**). In both cases, inhibitor treatment led to profound reduction of PI3K-driven DES1 activity – again with no major effects on DES1 protein. This extends our prior study (23) and shows that oncogenic PI3K-driven DES1 activation is connected to classical glucose metabolic pathways.

**Fig. 3.**
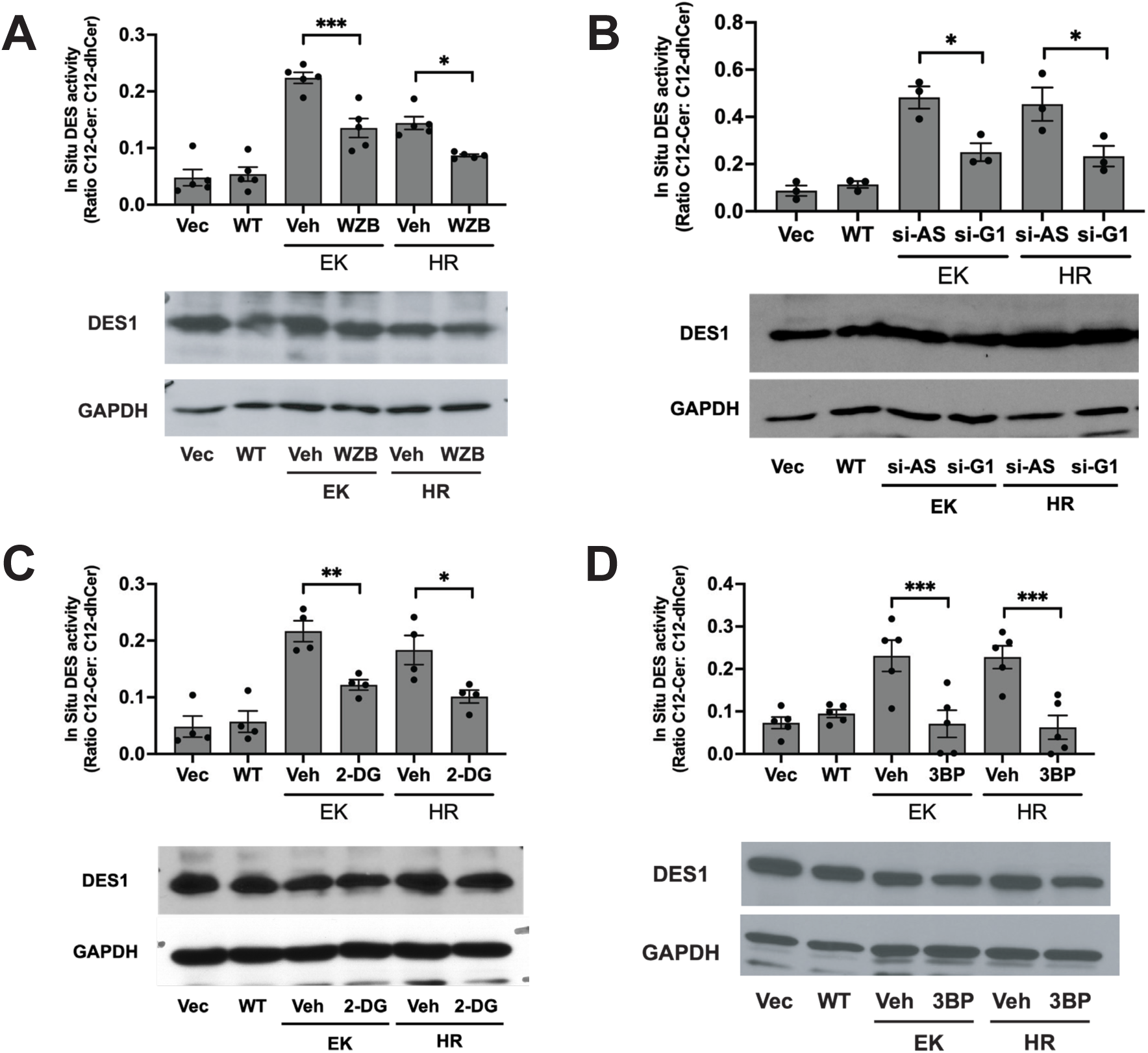
Oncogenic PI3K-driven DES1 activity requires glucose uptake and metabolism: 10A-PI3K cells were plated in suspension for 24h and in situ DES activity was assessed. a) Cells were treated with WZB (20mM) or vehicle for the course of the experiment; b) Cells were pre-treated with si-AS or si-GLUT1 (20nM) for 48h before being plated in suspension; c) Cells were treated with 2-DG (12mM) or vehicle; d) Cells were treated with 3-BP (… mM) or vehicle for the course of the experiment. For all experiments, protein was extracted from parallel treated samples and immunoblotted as shown. Data are expressed as mean ± SEM with * p<0.05, ** p<0.02, *** p<0.01 for comparisons shown.

### Oncogene-driven DES1 activity requires glucose flux through the pentose phosphate pathway (PPP)

Having consolidated a link between oncogenes, glucose metabolism, and DES1, it became important to understand which glucose pathways are connected to DES1 activation. The PPP is the first pathway to branch from glycolysis, has been linked to HER2- and SGK1-driven AIS (11, 38), and plays an important role in antioxidant defense through NADPH production. Given that NADPH and NADH function as major reducing agents for anabolic reactions including DES1 (39, 40), we assessed the requirement of the PPP for DES1 activity using the G6PD inhibitor (termed G6PDi, 20μM)(41) in both our 10A-NeuT (**Fig. 4A**) and 10A-PI3K cell lines (**Fig. 4B**). As in our prior results, all oncogenes were able to drive DES1 activity in suspension in the presence of vehicle, whereas G6PDi treatment led to a significant and comparable reduction in both NeuT (54%), PI3K-EK (58%) and PI3K-HR cells (53%) without effects on DES1 protein (**Supp Fig. 2A**). To consolidate this, siRNA was used to knockdown G6PD prior to culture in suspension. Immunoblot analysis confirmed siRNA efficacy in both cell types although as an intriguing aside, oncogenic NeuT was found to promote G6PD levels in suspension while oncogenic PI3K mutants had no effect on G6PD levels compared to 10A-Vec controls (**Fig. 4C**). Nonetheless, siG6PD led to significant inhibition of DES activity in 10A-NeuT (50%)(**Fig. 4D**) as well as both 10A-PI3K-EK (42%) and PI3K-HR cells (42%) (**Fig. 4E**) – similarly without effects on DES1 protein (**Supp. Fig 2B**). Taken together, this implicates glucose flux through the PPP as necessary for oncogene-driven DES1 activity.

**Fig. 4.**
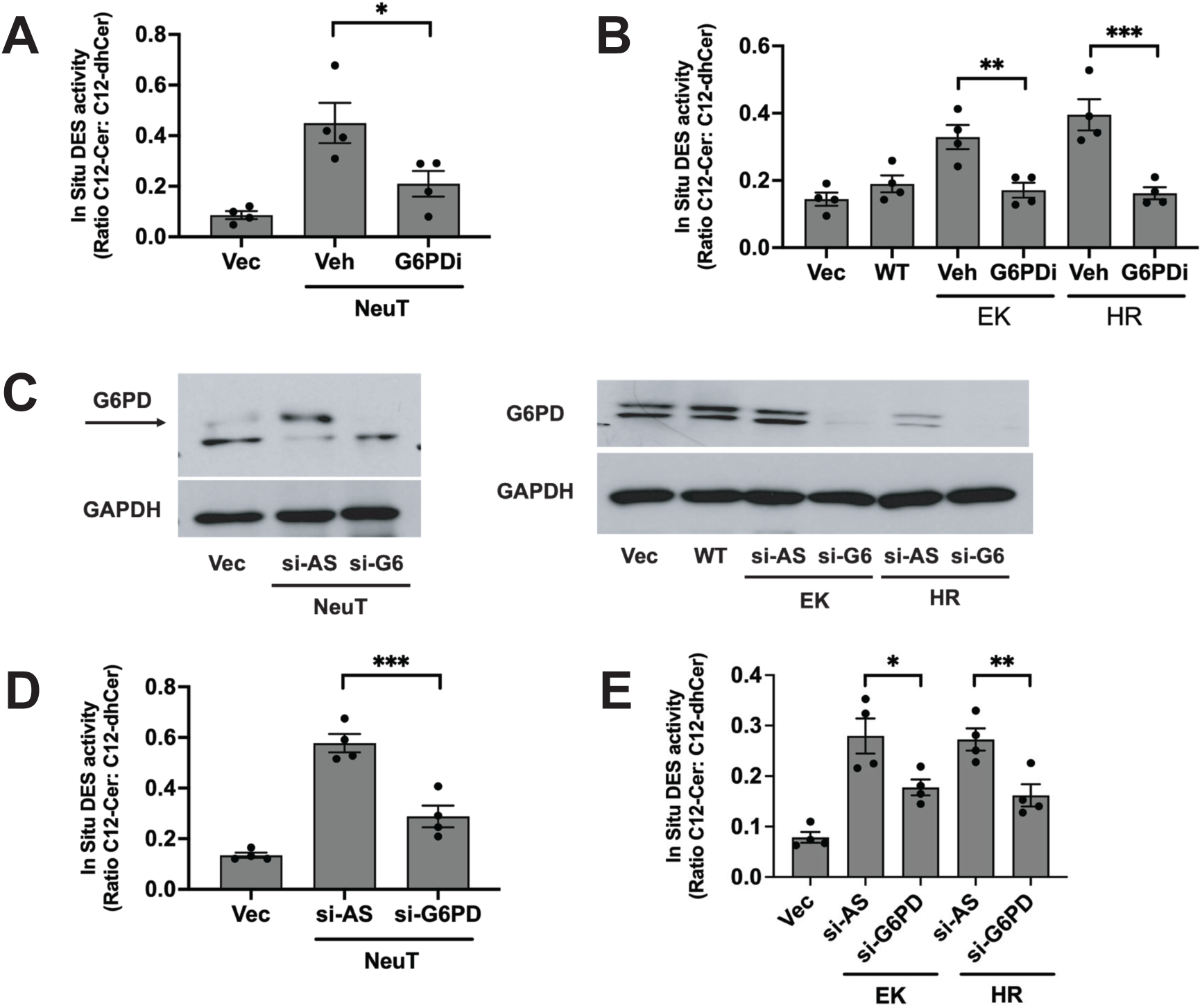
Targeting G6PD inhibits NeuT and PI3K-driven DES1 activity requires glucose uptake and metabolism: 10A-NeuT and 10A-PI3K cells were plated in suspension for 24h. a, b) Cells were treated with G6PDi (20uM) or vehicle and in situ DES activity was assessed; c) Cells were pre-treated with si-AS or si-G6PD (20nM) for 48h before being plated in suspension. Protein was extracted and immunoblotted as shown; d, e) Cells were treated as in ‘c’ and in situ DES activity was assessed. Data are expressed as mean ± SEM with * p<0.05, ** p<0.02, *** p<0.01 for comparisons shown.

### The PPP requirement for oncogene-driven DES1 activity is independent of antioxidant capacity

The PPP is a primary source of cellular antioxidants and upon ECM detachment, decreased PPP flux leads to oxidative stress (11). As increased oxidative stress can post-translationally inhibit DES activity (25), it was important to assess a role for ROS generation as a regulatory factor. For this, wild-type MCF10A cells were plated in suspension in the presence of the antioxidant N-acetylcysteine (NAC; 0.5mM) for 24h prior to in situ DES assays. Importantly, NAC was unable to reverse the decrease of DES activity seen in Susp (**Fig. 5A**) and had no effects on the levels of endogenous dhCer or Cer in MCF10A following ECM detachment (**Fig. 5B**). To consolidate this, 10A-NeuT cells were co-treated with G6PDi (20mM) and NAC (0.5mM) in Susp for 24h (**Fig. 5C**). However, as with wild-type MCF10A cells in suspension, NAC was unable to reverse the effects of G6PDi on DES1 activity suggesting the inhibition caused by G6PDi is not due to enhanced oxidative stress.

**Fig. 5.**
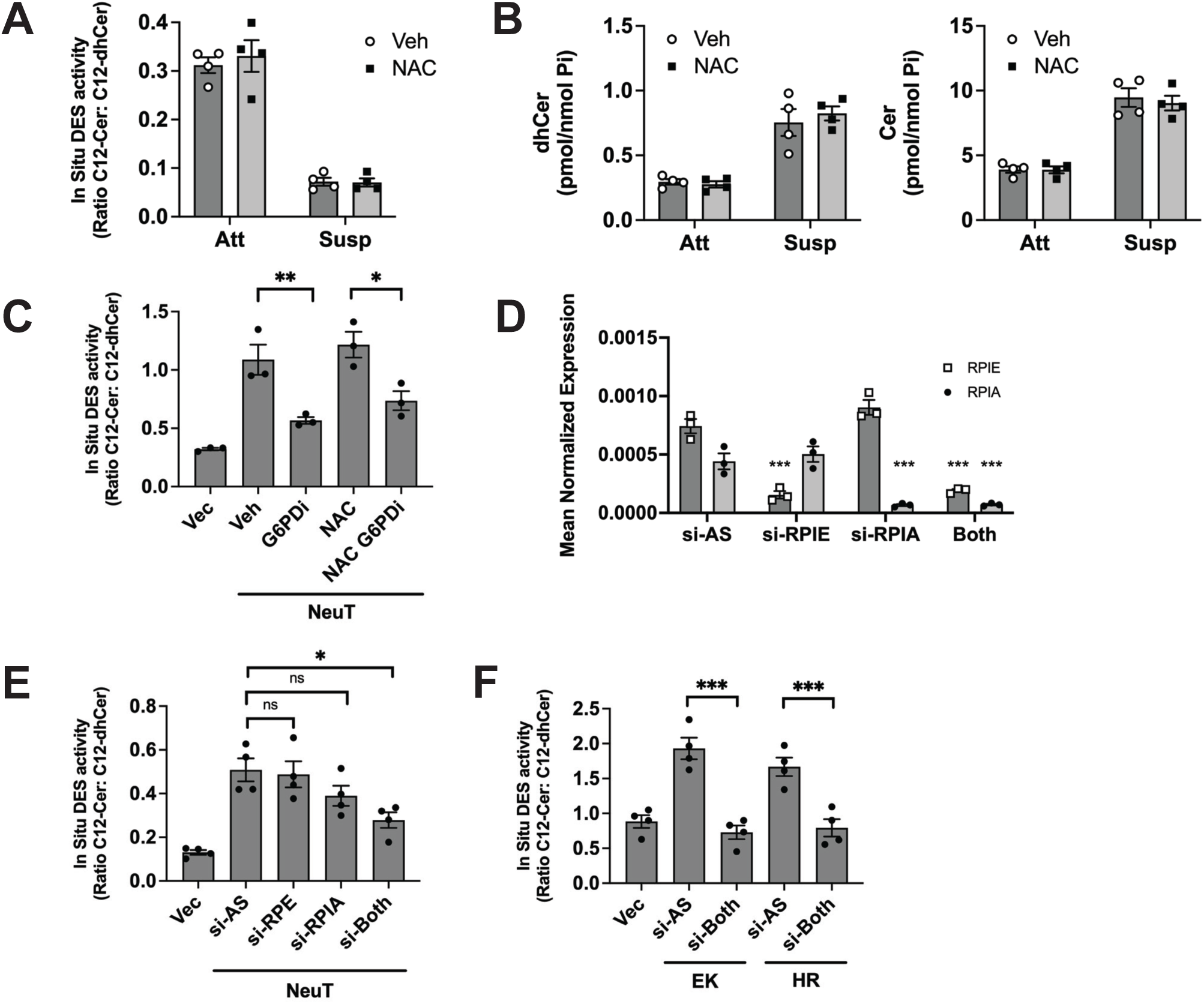
Oncogene-driven DES1 activity is independent of oxidative stress. a, b) MCF10A cells were plated in Att and Susp conditions for 24h with and without NAC (0.5mM). Cells were analyzed for *in situ* DES activity (a) and endogenous dhCer and Cer levels (b); 10A-NeuT cells were plated in suspension for 24h with vehicle, G6PDi (20uM), NAC (0.5mM) or both. In situ DES activity was assessed; d) Cells were pre-treated with si-AS, si-RPIE, si-RPIA, or both for 48h. Gene expression was assessed by qRT-PCR using actin as reference gene; e) 10A-NeuT cells were treated as in ‘d’ for 48h before being plated in suspension for 24h. In situ DES activity was assessed. f) 10A-PI3K cells were treated with si-RPIE and si-RPIA for 48h before being plated in suspension for 24h. In situ DES activity was assessed. Data are expressed as mean ± SEM with * p<0.05, ** p<0.02, *** p<0.01 for comparisons shown.

The PPP is divided into two phases – the non-reversible oxidative phase that leads to NADPH production and enhanced antioxidant capacity, and the reversible non-oxidative phase that is important for amino acid synthesis. The results above suggested a disconnect between the PPP requirement for DES activity and antioxidant capacity prompting us to assess if the non-oxidative phase of the PPP is also required for oncogene-driven DES1 activity. For this, siRNA against ribulose 5-phosphate 3-epimerase (RPIE) and ribulose 5-phosphate isomerase (RPIA) was utilized. As these enzymes can also compensate for each other, cells were treated individually and in combination with each siRNA. Real-time analysis confirmed efficacy of the siRNA for knockdown of each gene both when used in isolation and in combination (**Fig. 5D).** Analysis of DES activity in 10A-NeuT cells revealed that single knockdown of each gene had mild but insignificant effects while combined knockdown reduced DES activity to a level comparable to that seen with G6PD knockdown (**Fig. 5E**). A repeat of these studies in 10A-PI3K cells showed more striking effects with combined RPE/RPIA leading to almost complete inhibition of DES activity in both -EK and -HR cells. (**Fig. 5F**). Taken together, this shows that while the PPP is upstream of oncogene-driven DES1 activity, this does not appear to be related to its role in cellular antioxidant capacity and requires flux through the non-oxidative phase.

### Oncogene-driven DES1 activity requires glucose flux through glycolysis but not the TCA cycle

Recent studies have reported PI3K signaling influences glycolysis through regulation of aldolase (AldoA) activity (8). Given that PPP regulation of DES1 activity was independent of antioxidant capacity, and as the non-oxidative PPP rejoins glycolysis subsequent to AldoA, we speculated that metabolic regulation of DES1 may require combined glucose flux through glycolysis and the PPP. Accordingly, effects of AldoA knockdown on DES1 activity were assessed. Immunoblot analysis confirmed efficacy of siRNA at reducing AldoA protein levels with no effects on DES1 protein in either 10A-NeuT or -PI3K cells (**Fig. 6A**). Moreover, DES activity assays revealed that loss of AldoA led to significant decreases in DES1 activity in both 10A-NeuT (**Fig. 6B**) and 10A-PI3K cells (**Fig. 6C**). To determine if G6PD and AldoA were functioning in the same pathway upstream of DES1, combined G6PD and AldoA siRNA treatment was used in comparison to single siRNA treatments. In 10A-NeuT cells, both G6PD and AldoA siRNA (10nM) reduced NeuT-driven DES1 activity with a comparable level of inhibition for both. Moreover, combined knockdown of both did not have an appreciably larger effect (**Fig. 6D**). Similarly, in 10A-PI3K cells, combined AldoA and G6PD siRNA did not have additive effects on DES1 activity (**Fig. 6E** – only showing results with EK) although it should be noted that reducing the level of G6PD siRNA did not have as robust effects as seen before. Nonetheless, these results suggest that AldoA and G6PD function in the same upstream pathway of DES1 regulation.

**Fig. 6.**
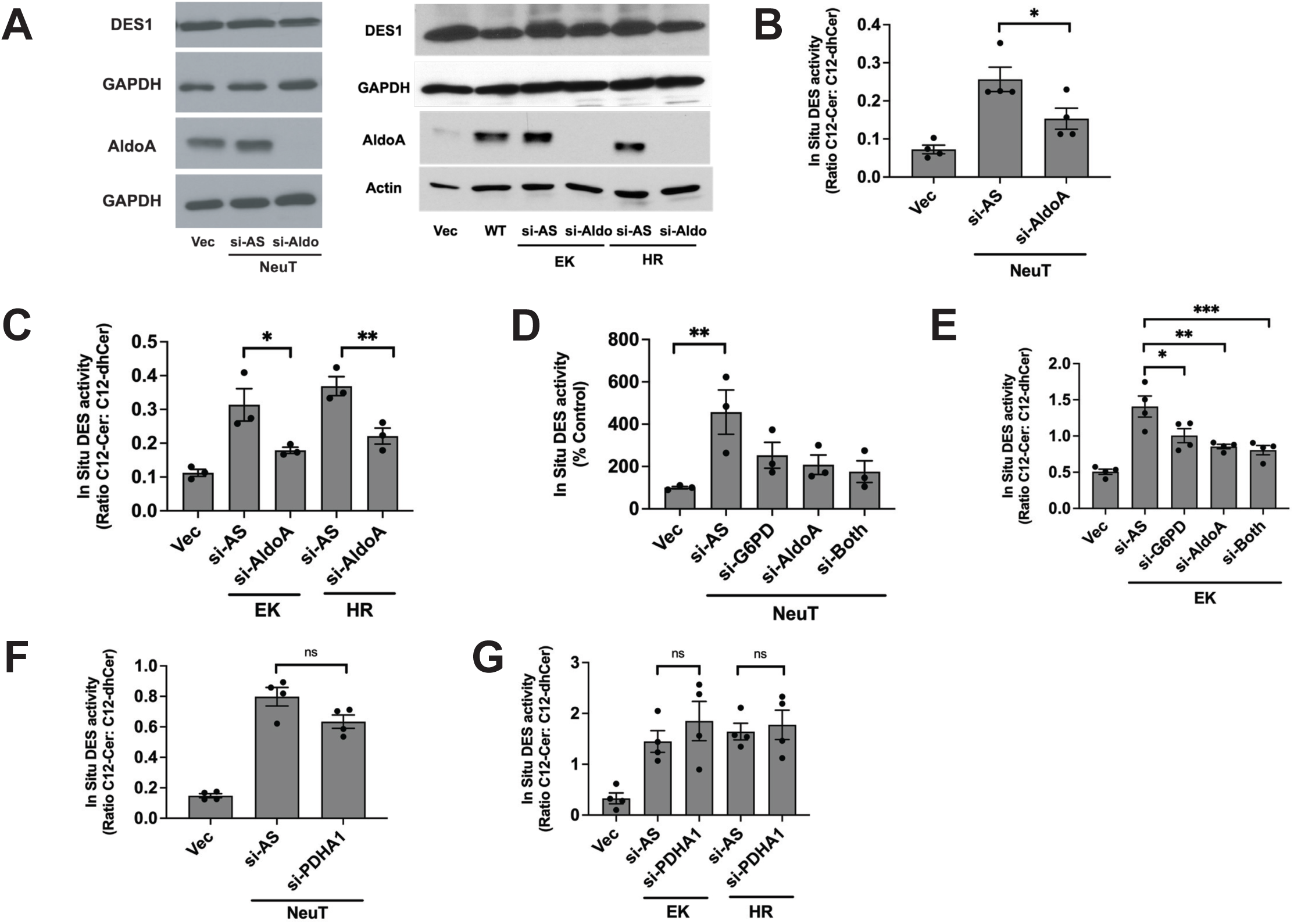
Oncogene-driven DES1 activity requires glucose flux through glycolysis but not the TCA cycle. a-c) 10A-NeuT and 10A-PI3K cells were pre-treated with si-AS or si-AldoA for 48h prior to being plated in susp for 24h. (a) protein was extracted and immunoblotted as shown. (b, c) In situ DES activity was assessed. (d, e) 10A-NeuT and 10A-PI3K cells were pre-treated with si-AS, si-G6PD, si-AldoA or both for 48h prior to being plated in susp for 24h. In situ DES activity was assessed. (f, g) 10A-NeuT and 10A-PI3K cells were pre-treated with si-AS or si-PDHA1 for 48h prior to being plated in susp for 24h. In situ DES activity was assessed. Data are expressed as mean ± SEM with * p<0.05, ** p<0.02, *** p<0.01 for comparisons shown.

The pyruvate produced from glycolysis can feed directly into the TCA cycle and onwards to oxidative phosphorylation in mitochondria to produce energy. As the metabolic regulation of DES1 occurs downstream of AldoA, it was important to determine if glucose flux into the TCA cycle is relevant – particularly as the TCA cycle is a source of cellular NADH and prior studies have linked the TCA cycle to AIS (42, 43). For this, we utilized siRNA against PDHA1, a key subunit of the pyruvate dehydrogenase complex that converts pyruvate into acetyl-CoA for the TCA cycle, with real-time PCR analysis confirming efficacy of the siRNA (**Supp Fig. 3**). In 10A-NeuT cells, PDHA1 siRNA had modest effects on DES1 activity (c. 20%) but this was not statistically significant (p >0.05; **Fig. 6F**). In contrast, PDHA1 siRNA had little to no effect on PI3K-driven DES1 activity (**Fig. 6G**). Taken together, this suggests that glucose flux into the TCA cycle is largely dispensable for metabolic regulation of DES1 driven by PI3K signaling in ECM-detached conditions.

### GAPDH is required for oncogene-driven DES1 activity

The non-oxidative PPP feeds back into glycolysis at two levels, with one such place being through glyceraldehyde 3-phosphate, which is subsequently metabolized by GAPDH. As data above suggested both PPP and glycolysis are linked with DES1 activity, this prompted us to assess if GAPDH was upstream of DES1 activity. Here, initial studies used koningic acid (KA) – also known as heptelidic acid – that is reported to directly inhibit GAPDH by binding to its covalent site (44, 45). For 10A-NeuT cells, treatment with 5uM KA led to a dramatic decrease in DES activity – reducing it to lower than that of Vec control cells (**Fig. 7A**) with similar marked effects seen in both 10A-EK and -HR cells (**Fig. 7B**). As with other intervention, this was without effects on DES1 protein (**Supp Fig. 4A**) consistent with post-translational regulation of DES1 activity. To consolidate these results, GAPDH siRNA was used although it should be noted that there were difficulties with obtaining effective siRNA downregulation of GAPDH. Indeed, 48h of pre-treatment was insufficient to get knockdown at the protein level (data not shown) and longer pre-treatment of 72h was used prior to plating in suspension – with effects on protein level often being modest compared to GAPDH expression (**Supp Fig. 4B**). Nonetheless, despite these challenges, modest GAPDH knockdown led to significant reductions in DES1 activity in 10A-NeuT (**Fig. 7C**) and 10A-PI3K cells (**Fig. 7D**) without effects on DES1 protein for either case. Taken together with the profound inhibitor results above, this suggests that GAPDH is a convergent point for post-translational metabolic regulation of DES1 driven by oncogenic activation of PI3K.

**Fig. 7.**
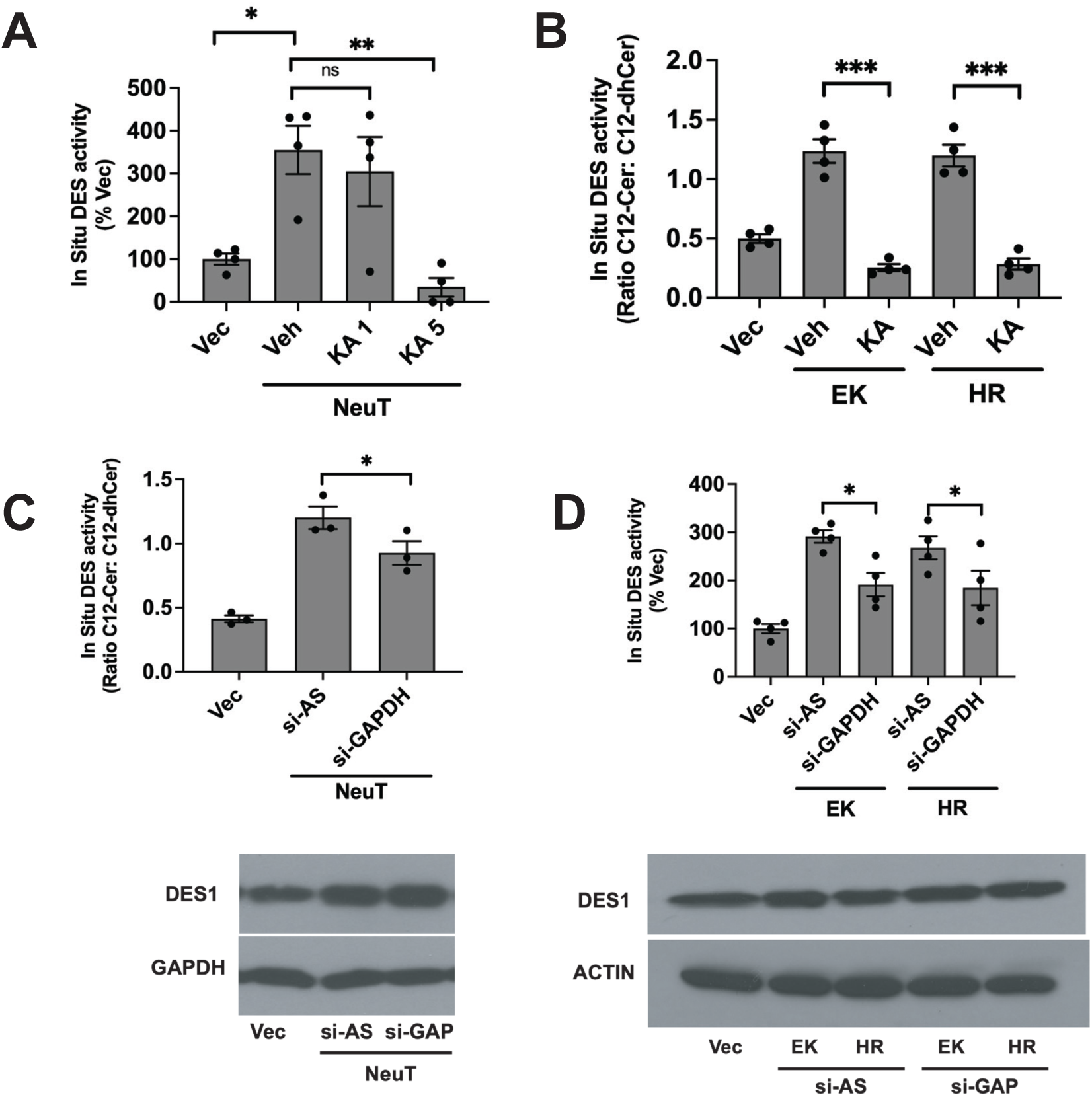
GAPDH is required for NeuT- and PI3K-driven DES1 activity. a, b) 10A-NeuT and 10A-PI3K cells were plated in Susp conditions for 24h with and without KA (1, 5uM). In situ DES activity was assessed; c, d) Cells were pre-treated with si-AS or si-GAPDH for 72h prior to being plated in susp for 24h. In situ DES activity was assessed. Data are expressed as mean ± SEM with * p<0.05, ** p<0.02, *** p<0.01 for comparisons shown.

## DISCUSSION

In comparison to other SL enzymes, little is known about the mechanisms of DES1 regulation which is particularly surprising given the fact that it directly controls the levels of two key bioactive SLs and is upstream of the synthesis of a host of other SLs including sphingomyelin and glucosylceramide. Our prior research established DES1 as a key HER2-regulated node in the SL network that was essential for the acquisition of AIS (23). Here, we have built upon this work, defining DES1 as a downstream effector of the oncogenic PI3K signaling pathway and establishing novel connections between DES1 and oncogene-driven glucose metabolism. This post-translational activation of DES1 requires glucose flux through both the PPP and glycolysis that converges on GAPDH. Taken together, this broadens the scope of DES1 activation in tumorigenesis to one of the most hyperactivated pathways in cancer and suggests wider potential for DES1 as a target of interest.

The deregulation of SL metabolism is well-established across many cancers (13–17) yet little is known about the underlying mechanisms that drive these changes both in terms of the key SL enzymes as well as the pathways or factors that drive their activation or inactivation. Here, our study significantly broadens the scope of DES1 as a key oncogene-regulated enzyme in the SL network and links its directly with hyperactivation of the PI3K pathway. Importantly, the effects of PI3K signaling on DES1 were observed with multiple mechanisms of activation including direct PI3K mutation, loss of tumor suppressors, and through other oncogenes i.e. K-Ras. This extends our prior study involving receptor-mediated PI3K activation (NeuT) and suggests that DES1 may have relevance across a wide range of cancers – particularly as increased PI3K signaling represents one of the most hyperactive signaling pathways in cancer [33, 34]. Biologically, this is consistent with our prior study establishing DES1 as a key mediator of AIS in BC [23] as PI3K has been widely reported to promote AIS and cancer aggressiveness in many cancers [**refs**]; indeed, we have also observed that siRNA knockdown of DES1 in 10A-PI3K cells effectively overcomes AIS and reduces colony formation (data not shown). However, given that PI3K has been implicated in other cancer-relevant biologies e.g. therapeutic resistance (46, 47), migration (48), and angiogenesis (49), it would be of interest to determine if DES1 activation is involved. Of note, while results connecting PI3K to DES1 were clear, results with other potential pathways were less so. Indeed, although activation of the ERK pathway with MEK-DD was not sufficient for DES1 activity in MCF10A cells, there was a modest effect of ERK inhibitors on DES1 activity in the context of oncogenic K-Ras. One possibility is that the ERK pathway functions as an additional regulatory pathway to ‘tune’ DES1 activity driven by PI3K signaling. Analysis of tumors primarily driven by hyperactive ERK signaling e.g. mutant RAF would shed further light on this and could establish important parameters for oncogenic activation of DES1. We also note that our pathway focus for this study was mostly on PI3K and ERK as these are commonly driven by a range of oncogenic signals. Thus, we also cannot exclude that involvement of additional oncogene-driven pathways e.g B-catenin, phospholipase C, protein kinase C in DES1 regulation.

The PI3K pathway functions as a key switch that connects nutrient availability with cell proliferation, growth, and survival and serves as a master regulator of glucose metabolism [3, 4]. Having previously linked NeuT-driven DES1 activity with glucose uptake [23], we here establish that glucose flux through both glycolysis and the PPP are necessary for NeuT and PI3K-driven DES activity, consistent with prior studies connecting PI3K with regulation of hexokinase [7], G6PD and the PPP [10, 11], and AldoA [8]. Notably, glucose flux into the TCA cycle was largely not necessary for oncogene-driven DES1 activity as targeting PDHA1 – that helps convert pyruvate into acetyl-CoA – had no significant effect. Of note, oncogenic regulation of PDHA1 was found to be dependent on the ERK pathway in ECM-detached conditions [42]. Thus, the lack of its requirement for NeuT-and PI3K-driven DES1 activity agrees with their ERK independence. Nonetheless, the requirement of both the PPP and glycolysis for DES activity suggests that the point of regulation does not occur within the confines of either pathway itself but more likely at a downstream convergence point.

Indeed, GAPDH – a point where the non-oxidative PPP can re-enter glycolysis – was necessary for oncogene-driven DES1 activity. Although this connection was previously unappreciated, this is consistent with prior reports suggesting GAPDH is a “guardian of the reductive capacity of the cells” although this was largely based upon its influence on NADPH generation and antioxidant capacity. Finally, while there were clear differences in the magnitude of effect of the inhibitor KA compared to siRNA, we do not believe this is a consequence of off-target effects. Indeed, KA has been reported to have high specificity for GAPDH [44, 45] and our studies were well within the reported range for robust inhibition. Alternatively, the more modest effects of siRNA could be due to our inability to get effective and sustained GAPDH knockdown – likely owing to its importance in cell metabolism and particularly in suspension when cells are undergoing metabolic stress due to ECM detachment. Nonetheless, in combination, these results establish GAPDH as an important upstream regulator of oncogene-driven DES1 activity.

Mechanistically, the metabolic regulation of DES1 occurs at the post-translational level – as DES1 levels were largely unchanged with all treatment. However, the nature of the mechanism connecting GAPDH and glucose metabolism with DES1 activity remains unclear. Although oxidative stress was shown to inhibit DES1 activity [25], results here suggest the metabolic regulation is independent of cellular antioxidant capacity. Thus, impacts of GAPDH on cellular NADH levels would also likely be irrelevant although it is possible that DES1 could be spatially coupled to a subcellular pool of NADH that is not broadly linked to antioxidant function. Alternatively, it is possible that specific metabolites downstream of GAPDH function as allosteric regulators of DES1 activation – akin to how ATP functions as an inhibitor of key respiratory enzymes [50, 51]. Given the broad number of metabolites and interlinked nature of glucose metabolism, investigations into this possibility are beyond the scope of the current paper but are the subjects of ongoing studies. Nonetheless, these findings have potential broader implications given known DES1 functions. For example, DES1 is widely established as a regulator of autophagy [52, 53] – thus, results here suggest that DES1 could have biological relevance in autophagy driven by low glucose. Similarly, DES1-driven Cer blocks translocation of GLUT4 to the plasma membrane through inhibition of Akt [54, 55]. Thus, results here suggest the possibility of a negative feedback loop whereby increased glucose uptake drives DES1 activity to ultimately reduce glucose transport. Finally, as oncogenes typically hijack normal signaling pathways within the cell, these results suggest the tantalizing possibility that DES1 functions as a point of crosstalk that helps co-ordinate SL synthesis with glucose availability In conclusion, this study sheds new light on mechanisms of DES1 regulation, establishing it as a downstream effector of the oncogenic PI3K signaling pathway and defining DES1 as a point of crosstalk connecting glucose and SL metabolism. In addition to advancing DES1 as a SL enzyme of interest in tumorigenesis, this suggests a potentially broader role for DES1 in co-ordinating SL synthesis with glucose availability. This could have implications for other DES1-relevant pathologies, particularly for metabolic diseases [56] and insulin resistance [57] where DES1 is a target of interest.

## ACKNOWLEDGEMENTS

The authors would like to acknowledge the technical support provided by the Cancer Center Lipidomics Shared Resource at Stony Brook University. We also would like to thank Zdislaw Szulc at the Lipidomics Synthetic Core of the Medical University of South Carolina for synthesis of the C12-dhCCPS substrate. We are also grateful to all members of the Lipid Cancer lab at Stony Brook University for helpful feedback and discussion throughout the duration of this study.

