## Supplemental Information for "Oncogene-driven metabolic regulation of Dihydroceramide Desaturase 1 (DES1) converges on GAPDH in matrix-detached conditions"

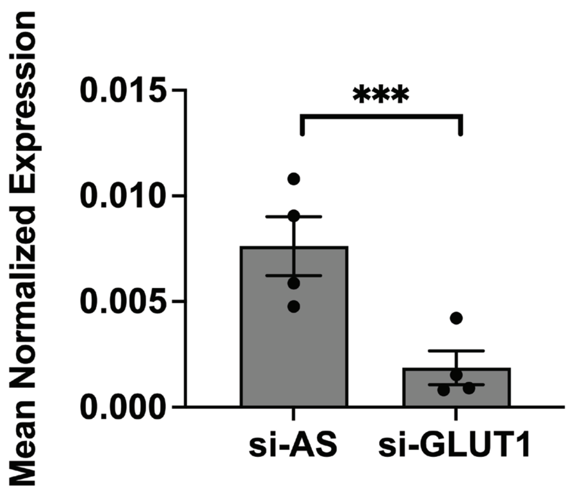


**Supp Figure 1**

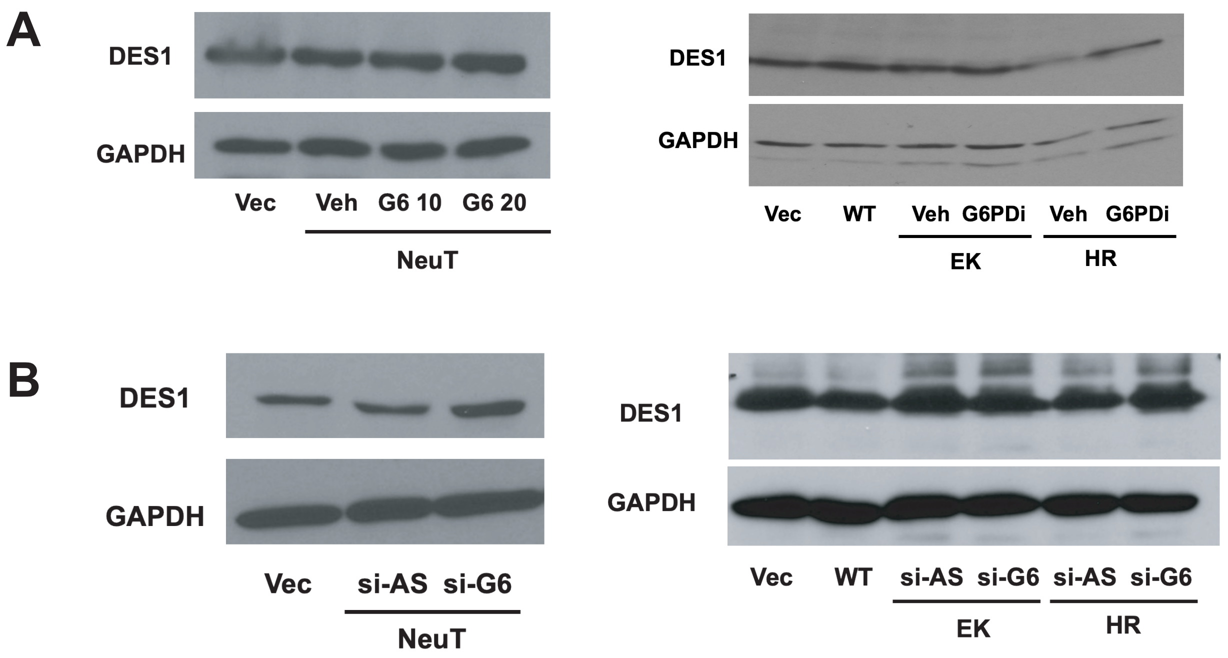


### Supp Figure 2

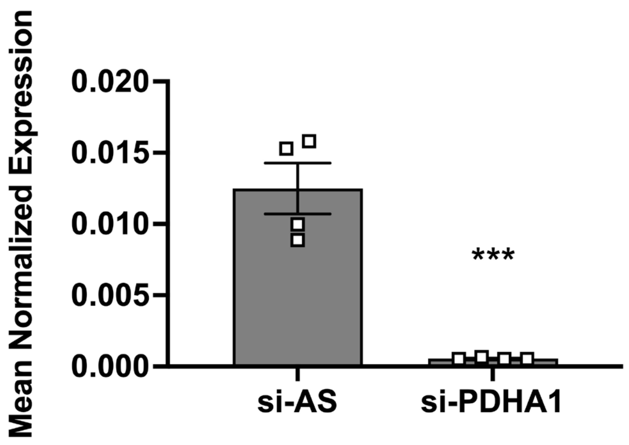


Supp Figure 3

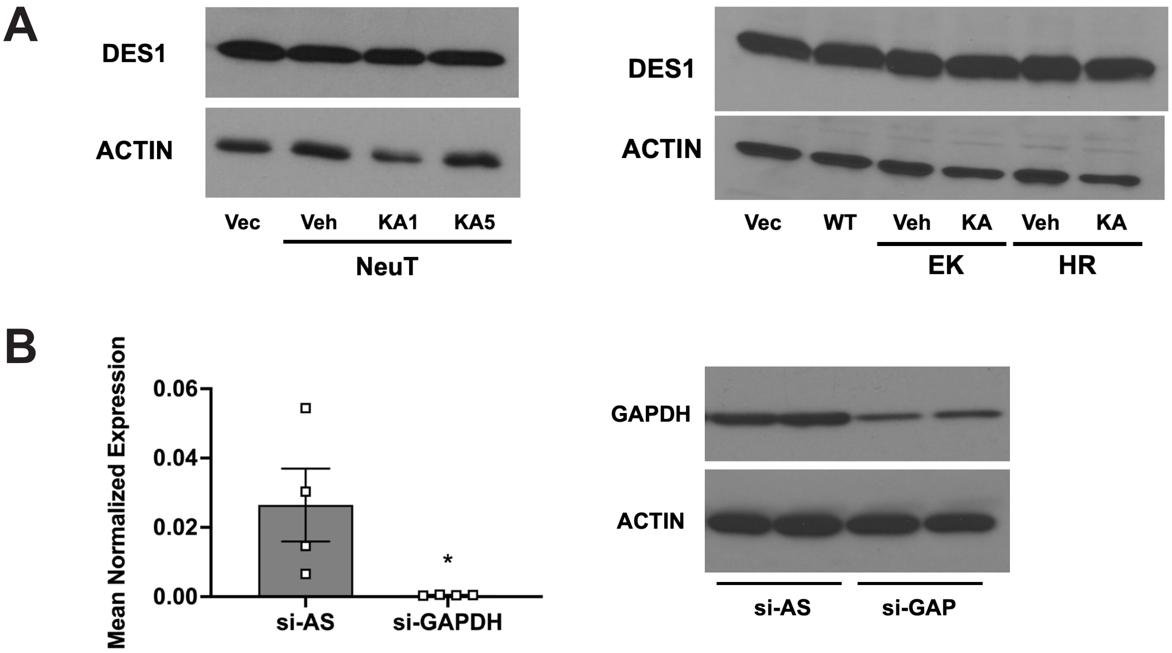


### Supp Figure 4

#### Supplemental Figure Legends

**Supplemental Figure 1. Validation of GLUT1 siRNA on GLUT1 expression**

10A cells were treated with negative control (si-AS, 20nM) or GLUT1 siRNA (20nM) for 48h. GLUT1 expression was analyzed by qRT-PCR with actin as reference gene.

**Supplemental Figure 2. Targeting G6PD with inhibitors and siRNA has no effect on DES1 protein levels**

(A) 10A-NeuT and 10A-PI3K cells were plated in suspension for 24h in the presence of vehicle or G6PDi. Protein was extracted and analyzed by immunoblot as shown.

(B) 10A-NeuT and 10A-PI3K cells were treated with si-AS (20nM) or si-G6PD (20nM) for 48h prior to being plated in suspension for 24h. Protein was extracted and analyzed by immunoblot as shown.

**Supplemental Figure 3. Validation of PDHA1 siRNA on PDHA1 expression**

10A cells were treated with negative control (si-AS, 20nM) or PDHA1 siRNA (20nM) for 48h. PDHA1 expression was analyzed by qRT-PCR with actin as reference gene.

**Supplemental Figure 4. Effects of KA on DES1 protein and GAPDH siRNA on GAPDH levels**

(A) 10A-NeuT and 10A-PI3K cells were plated in suspension for 24h in the presence of vehicle or KA as shown. Protein was extracted and analyzed by immunoblot as shown.

(B) 10A cells were treated with negative control (si-AS, 20nM) or GAPDH siRNA (siN1, 20nM) for 72h. GAPDH levels were analyzed by qRT-PCR (left) or immunoblot as shown (right).

| Gene Target | Company | Assay ID |
|---|---|---|
| AStar (negative) | Qiagen | 1027281 |
| PTEN | Qiagen | Hs_PTEN_6 |
| p85 | Life Technologies | s10535 |
| SLC2A1 (GLUT1) | Life Technologies | s12926 |
| G6PD | Life Technologies | s531165 |
| RPIE | Life Technologies | s224555 |
| RPIA | Life Technologies | s531630 |
| ALDOA | Life Technologies | s72 |
| PDHA1 | Life Technologies | s10243 |
| GAPDH | Life Technologies | s5572 |

**Supplemental Table 1: List of siRNA used**

| Target | Company | Catalogue # |
|---|---|---|
| p85 | Cell Signaling | 4257 |
| p110-alpha | Cell Signaling | 4255 |
| phospho-Akt (S473) | Cell Signaling | 4060 |
| Akt | Cell Signaling | 4691 |
| Actin | Sigma | A2228 |
| GAPDH | Cell Signaling | 5625 |
| phospho-ERK | Cell Signaling | 9101 |
| ERK | Cell Signaling | 4695 |
| DEGS1/MLD | Abcam | ab167169 |
| PTEN | Cell Signaling | 9188 |
| V5 | Life Technologies | R960-25 |
| G6PD | Cell Signaling | 8866 |
| Aldolase A | Cell Signaling | 8060 |

**Supplemental Table 2: List of antibodies used**

| Gene Target | Company | Assay ID |
|---|---|---|
| SLC2A1 (GLUT1) | Life Technologies | Hs00892681 |
| RPIE | Life Technologies | Hs00851816 |
| RPIA | Life Technologies | Hs01107136 |
| PDHA1 | Life Technologies | Hs04998681 |
| Actin (human) | Life Technologies | Hs01060665 |

**Supplemental Table 3: List of Taqman primers used**
